# CENP-A associated lncRNAs influence chromosome segregation in human cells

**DOI:** 10.1101/097956

**Authors:** Delphine Quénet, David Sturgill, Marin Olson, Yamini Dalal

**Affiliations:** Laboratory of Receptor Biology and Gene Expression, Center for Cancer Research; National Cancer Institute; National Institutes of Health; Bethesda, MD 20892, USA

## Abstract

Transcription occurs ubiquitously throughout non-coding parts of the genome, including at repetitive α-satellite DNA elements which comprise the majority of human centromeres. The function of temporally regulated centromeric transcription, and transcripts, is consequently a topic of intense investigation. In this study, we use high throughput approaches to identify and describe lncRNAs associated with the centromere specific histone variant CENP-A that arise from the transcription of specific centromeres at early G1, which we then show are physically associated with centromeres, and which are functionally necessary for accurate chromosome segregation. Targeted depletion of one such centromeric RNA, which originates from a single centromere, is sufficient to increase the frequency of chromosome segregation defects. These data support the emerging paradigm of the necessity of centromere-specific lncRNAs in the integrity of faithful chromosome segregation.

## INTRODUCTION

Centromeres are specialized chromosomal domains essential for kinetochore formation and attachment of the microtubule spindle, and consequently required for faithful chromosome segregation during cell division (1). The integrity of this chromatin region is maintained by a unique epigenetic landscape that is defined by the centromeric histone variant CENP-A, histone post-translational modifications, DNA methylation, chromatin-binding kinetochore proteins, and non-coding RNAs (2,3). Indeed, despite the lack of known genes in most centromeres, centromeric transcription is ubiquitous across eukaryotic species (4-19). In many of the species studied, the repetitive sequence of centromeres has contributed to the challenge in characterizing centromere-derived long non coding RNAs (lncRNAs). In humans, centromeric DNA consists of 171bp monomer α-satellite repeats (20). Arranged in a tandem fashion, these AT-rich repeats lack genes and share 40-100% sequence identity across different chromosome centromeres (21). The organization of monomeric or dimeric units of this base sequence into higher order repeats (HOR) can serve to distinguish individual centromeres (22). Recent work has demonstrated that lncRNAs are transcribed during late mitosis - early G1 by RNA polymerase II (RNAPII) from centromeric human α-satellite DNA, where they interact with the essential inner kinetochore proteins CENP-A and CENP-C (8,12,19). Excitingly, this phenomenon appears to be evolutionarily conserved across humans, Drosophila and Xenopus (17,18). In mammals, over-expression or down-regulation of centromeric α-satellite RNAs is associated with errors of chromosome segregation arising from centromeric defects during mitosis (10-12,19,23). Finally, a recent study in Drosophila confirms the importance of centromere transcription-assisted loading of CENP-A via factors such as FACT and Mis18α (24). The accumulation of evidence from multiple organisms synthesized into a working hypothesis suggests that centromeric transcripts and transcription are both essential for the accurate spatial and temporal targeting of centromeric proteins at the centromere, and consequently, contribute to accurate segregation of chromosomes during mitosis. Several outstanding questions remain to be answered: whether all centromeres in a given set of chromosomes are uniformly transcribed; the identity of these transcripts; whether centromeric transcripts have a discrete length reflecting a defined transcriptional unit; whether they bind *in cis* or *in trans* across different centromeres in a genome; the consequences of their loss or over-abundance; and finally, the mechanism by which cenRNAs might bind kinetochore proteins directly. In this study, we report our progress in investigating five out of these six important questions.

In our previous study (19), we reported that in early G1, RNAPII transcribes human centromeres, generating ~1.3kb long non-coding centromeric RNAs (cenRNAs) which are associated with soluble pre-assembly CENP-A/H4/HJURP‐ and CENP-A‐ chromatin-complexes. Depletion of these transcripts using α-satellite consensus sequence shRNAs leads to defective CENP-A loading in early G1 at a subset of centromeric fibers, resulting in chromosome mis-segregation and other cellular defects. These data, along with work of others cited above, strongly suggested a role for cenRNAs in maintaining the integrity of centromeric chromatin and cell division. In this study, to better characterize centromeric RNAs and their function, we performed soluble and chromatin bound CENPA associated RIP-seq (RNA-immunoprecipitation coupled to high throughput sequencing) from HeLa cells at early G1. Using this strategy, we identify several hundred cenRNAs which map to individual chromosomal centromeres, with strong homology to known centromeric α-satellite repeats, and which range from 300-2500 bases in length. Focusing our efforts on a subset of these cenRNAs, we show that one cenRNA, approximately ~1kb long, transcribes uniquely from chromosome 17, co-localizes with centromeric markers, and appears to be poly-adenylated. Functional characterization of this cenRNA by down-regulation of the transcript, demonstrates that its loss increases the frequency of chromosome segregation defects. These data support the emerging paradigm that centromeric RNAs are crucial players in the maintenance of centromeric integrity in higher eukaryotes.

## MATERIAL AND METHODS

### Tissue culture

HeLa cells were grown at 37°C, 5% CO_2_, in Dulbecco’s modified Eagle’s medium high in glucose and L-glutamine (#11965; Thermo Fisher Scientific/Gibco, Grand Island, NY) supplemented with 10% Fetal Bovine Serum (#26140-079; Thermo Fisher Scientific/Gibco) and 1X Pen/Strep solution (#10378-016; Thermo Fisher Scientific/Gibco).

Cells were synchronized by double thymidine block (0.5mM, #T9250; Sigma-Aldrich, Saint Louis, MO). After a first block of 19 hours, cells were released for 9 hours, followed by a second thymidine block of 16 hours. Then, cells were released for the appropriate time (10 hours for mitosis and 11 hours for early G1, as described previously in (19)).

### Antibodies

Antibodies are commercially available. Supplemental Table 1 lists all antibodies used for each experiment.

### shRNA

shRNAs incorporated into pGFP-V-RS plasmid vectors were purchased from Origene (Rockville, MD). Supplemental Table 2 lists sequences of these shRNA.

### Primers

Primers were purchased from Integrated DNA Technologies (Coralville, IA). Supplemental Table 3 lists primer sequence used for each experiment.

### BLAST on cenRNA#1

BLAST was performed on untranslated nucleotide sequence (blastn). Sensitive settings were: word size = 7 and expect value threshold = 10, with no low complexity filtering or repeat masking. Multiple blastn runs with variations of these parameters did not yield additional hits. To make our searching results more intuitive to interpret (as opposed to E-values and bit scores), a negative control sequence consisting of random sequence of the same length as cenRNA#1 (19) and with the same GC percent (47.4%, which is close to the human genome average of 46%; (25)) was generated. Random GC matched sequence for control was generated from the random DNA server at UC Riverside (http://www.faculty.ucr.edu/∼mmaduro/random.htm; Morris Maduro, accessed on February 2, 2015). The reference sequence of the 17bp CENP-B box (5’-CTTCGTTGGAAACGGGA-3’) is from (26). Adapter sequences identified in cenRNA#1 are described in (27).

### RNA Immuno-Precipitation – sequencing (RIP-seq)

RNAs associated to CENP-A or agarose bead (mock-IP) were purified and identified following the protocol described in (28). Briefly, for each condition, five F175 flasks of HeLa cells at a final confluency of 80% were used per RIP-seq. Trypsinized cells were washed two times with cold 1X PBS; 0.1% Tween 20 (#P2287, Sigma-Aldrich), before fixation with 0.1% formaldehyde (#15680, Electron Microscopy Sciences, Hatfield, PA) in 1X PBS. The reaction was stopped by addition of 125mM glycine (#50046, Sigma-Aldrich). Then, nuclei were isolated in TM2 buffer (20mM Tris– HCl, pH 8.0 (#15568-025, Thermo Fisher Scientific); 2mM MgCl_2_ (#AM9530G, Thermo Fisher Scientific); 0.5mM PMSF (#78830, Sigma-Aldrich); 1X complete protease inhibitor cocktail (#05892953001, Roche, Indianapolis, IN) complemented with 0.5% Nonidet^TM^ P40 Substitute (NP40, #74385, Sigma-Aldrich) and 10mM Ribonucleoside Vanadyl Complex (RVC, #1402, New England Biolabs—NEB, Ipswich, MA), and washed in TM2 buffer complemented with 10mM RVC. Chromatin was MNase-digested (MNase, #N3755, Sigma-Aldrich) in 0.1M TE buffer (0.1M NaCl (#24740-011, Thermo Fisher Scientific); 10mM Tris–HCl, pH 8.0; 0.2mM EGTA (#03777, Sigma-Aldrich) in presence of 2mM CaCl_2_ (#746495, Sigma-Aldrich). After addition of 10mM EGTA to stop the MNase action, nuclear pellet was resuspended in 1mL low-salt buffer (0.5X PBS, 5mM EGTA, 0.5mM PMSF, 1X complete protease inhibitor cocktail) complemented with 50 units of murine RNase inhibitor (#M0314S, NEB), and chromatin was extracted overnight at 4°C in an end-over-end rotator. After centrifugation, the supernatant was precleared with 30µL of protein A/G Plus agarose beads (#sc-2003,Santa Cruz Biotechnology, Dallas, TX) for 30 min at 4°C in an end-over-end rotator, and then, incubated with the anti-CENP-A primary antibody or no antibody for a minimum of 4 hours and a maximum of 12 hours at 4°C in an end-over-end rotator. CENP-A/primary antibody complex and mock-IP sample were immunoprecipitated with 50µL of protein A/G plus agarose beads for 2 hours at 4°C on the end-over-end rotator. Beads were washed three times with 1mL low-salt buffer containing murine RNase inhibitor and complete protease inhibitor cocktail. To isolate immuno-precipitated RNA, RNA–protein complex was eluted (elution buffer: 1% SDS; 0.1M sodium bicarbonate (#S6014, Sigma-Aldrich)), denaturated, treated sequentially with proteinase K (#AM2548, Thermo Fisher Scientific/Ambion) and DNaseI (#AM2222, Thermo Fisher Scientific/Ambion), and purified by phenol:chloroform:isoamylalcohol method. RNA concentration was determined by measurement on a UV-spectrophotometer and the quality of RNA was verified using a Bioanalyzer (#5067-1511, Agilent Technologies, Santa Clara, CA).

### RIP-seq library construction and sequencing

To prepare RIP samples for sequencing, libraries were constructed with the Illumina TruSeq ChIP Library Preparation Kit (IP-202-1012 & IP-202-1024, Illumina, San Diego, CA). RIP libraries were barcoded and run in multiplex on a HiSeq 2500 instrument (Illumina) by the NCI-Sequencing Core Facility. Cluster generation was performed using an Illumina HiSeq PE Cluster Kit v4 cBot (#PE-401-4001, Illumina). The sequencer was run for 125 cycles in paired-end mode. Base calling accuracy was high for each sample, with > 90% of bases with a quality score of at least Q30 (99.9% accuracy in Phred scale). RIP-Seq experiments were performed in duplicate, and included both, input and mock-IP samples as negative controls. Raw reads total for each sample ranged from 49 to 112 million reads.

### Computational analysis

To identify putative centromeric transcripts from the RIP-seq results, we followed a computational pipeline as described in our recently published methods book chapter on this topic (28). Briefly, raw reads in fastq format were cropped to remove low quality base calls from read ends with Trimmomatic (29). These pre-processed reads were then used in a first pass alignment to the repeating rDNA subunit (U13369.1, retrieved from GenBank) with Bowtie2 (30) in sensitive mode, to deplete ribosomal transcripts *in silico*. The unaligned reads from this process were used for further analysis. These reads were mapped to the Build 38 reference genome (hg38) using Tophat v2.1.0 (31). For Tophat alignment, up to three read mismatches (and an edit distance of three) were allowed for alignment, and only a single best alignment was reported for each read (the ‘-g 1’ option). The centromeric sequence included in this version of the reference allowed us to map reads deriving from centromeric transcripts. Transcripts were defined from these alignments by reference-guided transcript assembly using Cufflinks (31). These assemblies were merged and transcript abundances were estimated with Cufflinks. Quantitative tracks for browser visualization were generated using deepTools (32). For visualization, Bowtie2 alignments were run in local mode (which allows soft-clipping), and enrichment relative to input was plotted as the subtraction of the depth normalized coverage in the input sample from the depth normalized coverage in the IP sample.

Open reading frame (ORF) finding was performed using the NCBI Orf Finder tool (https://www.ncbi.nlm.nih.gov/orffinder/), selecting “ATG only” to define ORF start sites. To search for TATA-boxes, we used the “fuzznuc” Emboss package that can map the consensus motif with amibiguity codes (“TATAWAAR”; (33)). For evidence of other gene elements, we used Promoter 2.0 and Genscan on the region +/- 2kb from the cenRNAs (34,35).

Motif enrichment was performed with MEME (36). Since this tool limits the amount of sequence that may be input at one time, we separated out transcript sequences into four subsets and ran them individually. The maximum motif size was set to 171bp, the length of the α-satellite monomer. We analyzed the ten highest reported scoring motifs. Sequence logos for the top-scoring motif in each subset is presented in Supplemental Figure 1A. Each of these motifs were highly similar to α-satellite monomer (88-98% identical over 98-100% of the query).

To generate distance trees for the complete set of cenRNAs, we aligned them using MUSCLE (Multiple Sequence Comparison by Log-Expectation; (37)), and calculated a tree based on average distance using percent identity (Supplemental Figure 1B). These calculations were implemented with Jalview (38). The results were then visualized in Dendroscope 3 to make the tree more interpretable (39). Since branching patterns can differ by algorithm, we generated multiple trees for comparison. Representative of this analysis, trees using alternative alignments Clustal and clustering performed with Unweighted Pair Group Method with Arithmetic Mean (UPGMA), including bootstrap results (using Geneious v.10.0.6; (40)), are shown in Supplemental Figure 1C-F. We note that a fraction of transcripts do not cluster at the bootstrap level of 70% support, but despite this, separation of transcripts into clusters was observed consistently.

### Poly-adenylation of candidate cenRNAs

Total RNAs from HeLa cells were purified with Trizol reagent (#15596026, Thermo Fisher Scientific/Ambion) following manufacturer instructions. To run out genomic DNA contamination, these total RNAs were incubated with 4 units of DNase I (#M0303S, NEB) for 30 min at 37°C. The reaction was stopped by addition of 50mM EDTA (#351-027-721, Quality Biological, Gaithersburg, MD) and incubated at 75°C for 10 min. Then, total RNAs were purified using the Zymo Clean & Concentratator kit (#R1015, Zymo Research, Irvine, CA) before being retro-transcribed using SuperScript IV kit (#18091050, Thermo Fisher Scientific) with either polydT primers or random hexamers following manufacturer instructions. PCR reaction was run using Dream Taq PCR master mix kit (#K1071, Thermo Fisher Scientific) with α-satellite or cenRNA#4 primers (Supplemental Table 3). Cycling conditions for PCR were: 30s at 98°C; 30 cycles: 5s at 98°C, 30s at 57°C, 20s at 72°C; 300s at 72°C. Finally, PCR products were analyzed on 2% agarose gel.

### RNA fluorescence *in situ* hybridization (RNA FISH)

Custom Stellaris RNA FISH probes labeled with Quasar dyes (*i.e.*, Quasar^®^570 or Quasar^®^670) were designed against specific cenRNAs and purchased from Biosearch Technologies (Petulama, CA). 75% confluent HeLa cells on poly-L lysine coverslip in 6-well plate were washed three times with HANKS buffer (#14170112, Thermo Fisher Scientific) and fixed with 4% paraformaldehyde (#15714, Electron Microscopy Sciences) in 1X PBS for 10 min at room temperature (RT). After washing cells with 1X PBS two times for 10 min, cells were made permeable in 70% ethanol (#61509-0010, Acros Organics, Pittsburgh, PA) overnight at 4°C. In order to confirm that the observed RNA FISH signal results from the hybridization of the probe with an RNA molecule and not genomic DNA, for every RNA FISH experiment, cells were rinsed once with 1X PBS in presence or absence of 1mg/mL of RNase A (#12091-021, Thermo Fisher Scientific). Cells were pre-incubated with 2X SSC (#AM9763, Thermo Fisher Scientific/Ambion); 10% deionized formamide (#15745, Electron Microscopy Sciences) for 5 min, and incubated with hybridization mix (0.1µM RNA FISH probe set diluted in 10% dextran sulfate (#S4030, EMD Millipore, Billerica, MA), 2X SSC, 10% deionized formamide), for 4 hours to overnight at 37°C in the dark. Finally, cells were washed twice with 10% deionized formamide in 2X SSC for 30 min at 37°C and once with 2X SSC for 5 min at RT. Coverslips were mounted on cells with anti-fade mounting medium Prolong Gold with DAPI (#P36935, Thermo Fisher Scientific).

### Immuno-fluorescence / RNA FISH (IF/RNA FISH)

IF/RNA FISH experiments were performed as described previously (28). Custom Stellaris RNA FISH probes were purchased at Biosearch Technologies. Briefly, 75% confluent cells on poly-L lysine coverslip in 6-well plate were washed three times with HANKS buffer and fixed with 4% paraformaldehyde in 1X PBS for 10 min at RT. After permeabilization in 0.1% Triton X-100 (#T8787, Sigma-Aldrich) in 1X PBS for 5 min at RT, cells were incubated with anti-CENP-B primary antibody in IF buffer (1X PBS; 1% normal goat serum (#005-000-121, Jackson ImmunoResearch, West Grove, PA); 50 units of murine RNase inhibitor) overnight at 4°C. After three washes in 1X PBS, cells were incubated with secondary antibody (goat anti-rabbit IgG (H+L) secondary antibodies, Alexa Fluor®488 or Alexa Fluor®568 conjugate (Thermo Fisher Scientific) in IF buffer for 1 hour at RT in the dark. To validate the RNA signal, half of slides were treated with 1X PBS complemented with 1mg/mL of RNase A. Cells were fixed with 4% paraformaldehyde in 1X PBS for 10 min, and washed twice with 1X PBS for 10 min. Then, cells were treated for RNA FISH as described earlier in “RNA FISH” section.

### Electroporation of HeLa cells - shRNA depletion of centromeric RNAs

1.2×10^6^ cells were mixed with 100µL of RT nucleofection reagent (#VACA-1001, Lonza, Allendale, NJ) and 7µL of plasmid containing shRNA, or a GFP control to quantify transfection efficiency (500ng/µL of Scrambled, cenRNA#4A, cenRNA#4B, α-satellite A, α-satellite B, H3-GFP). After electroporation with Nucleofector^®^ Device (Lonza, Walkersville, MD) using program O-005 (for high viability), HeLa cells were resuspended in 500µl of warm media. 200µL of cells were transferred to a 6-well plate. Two days later, cells were stained for RNA FISH to validate the depletion in cenRNA. Otherwise, they were selected with 0.5µg/mL puromycin until day 6, at which they were fixed and treated for IF.

### Immuno-fluorescence

HeLa cells were grown on poly-D-Lysine-treated coverslips in 6-well plate. After two washes with cold 1X PBS, cells were prefixed for 30s with cold 4% paraformaldehyde in PEM (80mM K-PIPES pH6.8; 5mM EGTA pH7; 0,2mM MgCl_2_). After washing three times with cold PEM, soluble proteins were extracted for 5 min on ice with 0.5% Triton X-100 in CSK (10mM K-PIPES pH6.8, 100mM NaCl, 300mM sucrose, 1mM EGTA, 3mM MgCl_2_). Few drops of 4% paraformaldehyde in PEM were added for 5 min. Coverslips were then incubated with fresh 4% paraformaldehyde in PEM for 40 min on ice. After three washes with PEM, cells were permeabilized with 0.5% Triton X-100 in PEM for 30 min at RT, washed again three times, and blocked in 1X TBS, 3% Bovine Serum Albumin, 5% normal goat serum for 1 hour at RT. Finally, cells were incubated with the primary antibody diluted in 1X TBS, 1% Bovine Serum Albumin, 5% normal goat serum overnight at 4°C in a humidified chamber. Cells were washed three times for 5 min at RT with 1X TBS, 0.1% Tween 20, and incubated with secondary antibody for 1 hour at RT. After washing, coverslips were mounted on slides with anti-fade mounting medium Prolong Gold with DAPI.

### Microscopy observation and analysis

RNA FISH, IF and IF/RNA FISH slides were observed with a DeltaVision or DeltaVision Elite RT microscopy imaging system (GE Healthcare, Pittsburgh, PA) controlling an interline charge-coupled device camera (Coolsnap, Tucson, AZ) mounted on an inverted microscope (IX-70; Olympus, Center Valley, PA). Images were captured by using a 60X objective at 0.2 µm z-sections and analyzed with Image J (1.50e; Java 1.6.0_20, NIH, Bethesda, MD).

### Statistical analysis

On each figure is indicated the number of repetition for each experiment (generally N=3 biological replicates; typically, a minimum of 100 cells - n=100 - scored per phenotype measured). Standard error was determined for all quantification measures. To test the significance of chromosome segregation defect measurements, a two-tailed Fisher’s exact test was performed. For all tests, α was assumed to be 0.05. Significant p-values are indicated on the figures or tables with an asterisk each time it was evaluated.

## RESULTS

### Identification of centromeric RNAs by RIP-seq originating from discrete centromeric satellite sequences

To design an unbiased approach to identify potentially repetitive centromeric RNA sequences, we constructed libraries from RIP experiments followed by high-throughput sequencing (RIP-seq), and identified several hundred RNA species associated with CENP-A at centromeres (Figure 1A; (28)). We refined a computational pipeline to assemble putative centromere-derived transcripts from RIP-seq reads (Figure 1B; (28)). Briefly, reads deriving from ribosomal transcripts were depleted *in silico,* mapped to Build 38 reference sequence including centromere sequence models, and assembled into putative transcripts using the Build 38 reference as a guide. This analysis yielded 432 putative transcripts that were at least 300bp in length (the approximate library fragment length), with abundance > 1 fragment per kilobase per million mapped reads (FPKM) in the chromatin fraction of each of two replicates, and which mapped to centromeres (Figure 1C-F and Supplemental Table 4). The complete set of candidate cenRNAs has a median size of 443bp, and a maximum size of 2450bp (Figure 1G). They were detected on every chromosome except for chromosomes 6, 13, and 14, although low abundance transcript predictions were defined on these chromosomes as well (Figure 1H). Most displayed reduced enrichment in shRNA α-satellite samples (60-66% of cenRNAs). These results demonstrate the pervasive nature of centromeric transcription in the HeLa transcriptome, and suggest that centromeric transcripts span a size range from 300-2500 bases, arising from transcripton of a large fraction of centromeres in the human nucleus.

We next searched for recurring motifs in these centromeric RNA sequences using the algorithm MEME. Reassuringly, the top scoring motifs were highly similar to the canonical α-satellite monomer (Supplemental Figure 1A), confirming the predominance of α-satellite sequence in these transcripts. To analyze the sequence heterogeneity in our putative cenRNAs, we generated trees computed on percent identity distance (Supplemental Figure 1B-F). Although α-satellite sequence is predominant in these transcripts, sufficient differences exist to separate them into four distinct groups, or ‘clades.’ Although transcripts from the same centromere tended to group together, the branching was not strictly by chromosomal origin. These results demonstrate the diversity of centromeric transcribed sequences, and show that they are separable and identifiable in sequencing experiments.

From the RIP-seq centromere-mapping putative transcripts identified above, we selected three candidate cenRNAs for follow-up experiments, based on relative abundance and enrichment relative to controls (cenRNA#2, cenRNA#3, cenRNA#4; Figure 1C-E). Interestingly, each selected cenRNA was also detected in soluble pre-assembly CENP-A RIP-seq, but at less than half the abundance, supporting a relatively stable chromatin association for these targets. A reduction of enrichment was also observed in RIP samples transfected with α-satellite shRNA, suggesting these cenRNAs are depleted in our knockdown experiments (8.4-79% for cenRNA#2, 24.9-31.2% for cenRNA#3, and 9.2-31.7% for cenRNA#4; Supplemental Table 4). Each of these transcripts was defined by alignment to centromeric reference assembly, containing sequence in previously described α-satellite monomers (20). Using this approach, we observed that cenRNA#2 maps to chromosome 3 with high sequence similarity to the HOR DSZ1 (41), although RepeatMasker classifies the majority of the transcript region as scaffold attachment region (SAR) class satellite rather than ALR/alpha (42,43). In contrast, cenRNA#3 and cenRNA#4 mapped with very high homology to centromeric sequences on chromosome 17, specifically to the D17Z1 α-satellite array (44,45). Thus, these results show the existence of multiple centromeric long non-coding RNAs originating from several chromosomes. In addition, multiple RNAs can be transcribed from one chromosome. We interpret these data to indicate two points. First, that individual centromeres from different chromosomes are competent for transcription, and second, that transcription likely does not have a distinctive single start site per centromere.

We analyzed the sequence characteristics of the cenRNAs for evidence of coding potential. None of them included an open-reading frame (ORF) with 100 or more codons (a commonly used cut-off for defining a lncRNA; (46)). We also searched for protein alignments to six-frame translated cenRNA sequence. Neither cenRNA had evident similarity to human protein, with best hits all from predicted microbial protein sequence (Supplemental Datafile 1). This analysis supports the commonly accepted notion that centromeric transcripts are non-coding.

### Centromeric RNAs are present at discrete centromeric loci in the nucleus during late mitosis-early G1

We, and others, have previously shown that human centromeres are preferentially transcribed by RNAPII at late mitosis – early G1 (8,19). To assess the abundance of expression of these newly-identified cenRNAs in human cells at early G1, we synchronized HeLa cells to that specific time of the cell cycle and performed RNA FISH. Then, we measured the frequency of cells expressing these cenRNAs by counting cells with a bright RNA FISH focus consistent with known appearance of transcriptionally active foci.

For RNA FISH localization controls, we chose MALAT1 and NEAT1, two well described nuclear lncRNAs, which are ubiquitously expressed in human cells (47). MALAT1 was highly expressed with punctated spots all over the nucleus, whereas NEAT1 signal was more discrete with an average of 7 spots per cell (Figure 2A). This distribution is consistent with the localization of MALAT1 and NEAT1 described in the literature (47), implicating our approach conserve the spatial localization of lncRNAs. Both MALAT1 and NEAT1 were expressed in more than 80% of late mitosis – early G1 synchronized cells (Figure 2B). To exclude an unspecific hybridization of the RNA probes to genomic DNA, cells were treated with RNase A. As expected, the RNA FISH signal was diminished substantially after RNase A treatment, confirming the the ribonucleic nature of the observed signal and validating the method (Supplemental Figure 2).

Next, we studied the localization of centromeric α-satellite RNAs (Figure 2). Consensus centromeric transcripts were revealed by probes designed against the consensus sequence of α-satellite repeats derived from (20). On average, seven discrete nuclear spots were observed in 34% of late mitosis – early G1 synchronized cells, but none in cells treated with RNase A (Figure 2B and Supplemental Figure 2). This result suggests that several centromeric RNAs are transcribed from multiple centromeres and are localized in multiple nuclear localizations at the end of mitosis.

Next, we analyzed the localization of the three selected cenRNAs, namely cenRNA#2, cenRNA#3 and cenRNA#4 (Figure 1) using the same approach. Using probes designed against cenRNA#2, we observed RNA FISH signals were diffuse and similar to background and the RNase A-treated control, suggesting these probes were prone to non-specific recognition of other RNAs in the genome (Figure 2A and Supplemental Figure 2). In contrast, RNA FISH experiments performed with RNA probes designed against cenRNA#3 and cenRNA#4 expressed well defined and unique foci in synchronized HeLa cells (Figure 2A). cenRNA#3 and cenRNA#4 were observed in ∼28% and ∼37% of late mitosis - early G1 synchronized cells, respectively (Figure 2B). These signals were lost upon RNase A treatment, confirming the ribonucleic nature of the analyzed foci (Supplemental Figure 2).

Next, we characterized these transcripts by determining the chromatin domain to which they are localized. For this purpose, we performed a co-IF/RNA FISH to observe their relative localization to centromeres, using CENP-B, which marks centromeric DNA (Figure 3A). Since these RNAs were originally identified by CENP-A-chromatin RIP-seq (Figure 1), we expected to observe a colocalization or partial co-localization of cenRNAs with CENP-B domains. Reassuringly, in cells expressing centromeric α-satellite transcripts, ∼74% of them displayed overlap between CENP-B and α-satellite RNA signals (Figure 3B). The absence of RNA FISH signal after RNase A treatment confirm that the observed α-satellite foci were indeed centromeric transcripts and not genomic sequence (Figure 3B). Next, we assessed the relative localization of cenRNA#4 to centromeres by repeating this co-IF/RNA FISH experiment (Figure 3C). As observed for α-satellite transcripts, cenRNA#4 was also found overlapping with CENP-B signal in ∼70% of cells expressing cenRNA#4 (Figure 3C). Upon RNase A treatment, the RNA signal was lost (Figure 3C). Thus, our results indicate that in late mitosis - early G1 cells, cenRNA#4 and α-satellite transcripts localize adjacent to CENP-B domains.

### Centromeric transcripts are poly-adenylated

Since centromeric RNAs have been reported to be polyadenylated in Drosophila (18), we were curious whether human centromeric transcripts might also contain a poly(A) tail. To test this hypothesis, we retro-transcribed DNase I-treated total human RNAs with either poly(dT) or random hexamer primers, amplified cDNAs and compared profiles of PCR products on agarose gel (Supplemental Figure 3). Using primers designed against centromeric consensus sequence (α-satellite), we observed discrete bands every ∼170 bases (from 170 to 680 bases) after retro-transcription with both poly(dT) or random hexamer primers (Supplemental Figure 3A). These data suggest that these centromeric transcripts are likely polyadenylated. To confirm the ribonucleic nature of the analyzed molecules, several controls were performed. To prove the absence of contaminant in our reaction mix, the experiment was performed without total human RNAs (-RT in Supplemental Figure 3A). We did not observe PCR products in this condition. We also tested the property of primers to self-anneal or dimerize by running a PCR reaction without cDNA (-cDNA in Supplemental Figure 3A). No PCR products were amplified. Altogether, these data suggest that centromeric RNAs are polyadenylated. We decided to further test this hypothesis that cenRNAs present a poly(A) tail by repeating the experiment with primer sets designed against cenRNA#4 (Supplemental Figure 3B).Similarly to centromeric α-satellite transcripts, several discrete bands were observed every ∼170 bases after retro-transcription with either poly(dT) or random hexamer primers (Supplemental Figure 3B). However, when total RNAs were retro-transcribed, an additional band with an approximate size of 50 bases was amplified. The same band is observed in control conditions (-RT, and –cDNA; Supplemental Figure 3B). The difference between profiles between the assay (poly(dT) and random hexamers) and controls (-RT, and –cDNA) indicated that cenRNA#4 is poly-adenylated as well (Supplemental Figure 3B).

It has recently been reported in Xenopus (17), that centromeric RNAs appear to be spliced before they function to assist in CENP-A/CENP-C loading. We attempted to investigate whether human cenRNAs are processed using a 3’-5’ RACE experiment; however, these results were inconclusive. Thus, while we were unable to definitively conclude whether human centromeric RNAs undergo splicing, these data suggest that polyadenylation is a maturation step that cenRNAs undergo before they bind CENPA/HJURP complexes.

### Down-regulation of centromeric RNAs leads to chromosome segregation defects

In our earlier work, the down-regulation of consensus α-satellite RNAs by shRNA treatment led to an accumulation of mitotic defects (19). Indeed, parallel and previous work in Drosophila, Xenopus, and mammals demonstrate that loss of centromeric RNAs leads to mitotic defects underpine by the loss of CENP-A and CENP-C (15,17-19,48). Here, in order to investigate the function of cenRNA#4, we generated an shRNA construct designed against cenRNA#4 and analyzed the outcomes of the down-regulation of this transcript (Figure 4).

HeLa cells were transfected with the H3-GFP or shRNA constructs and treated with puromycin to select for transfected cells (Figure 4A). At Day 2, the efficiency of transfection was assessed by quantifying the percentage of cells expressing H3-GFP and by an RNA FISH approach (Supplemental Figure 4). The percentage of cells expressing α-satellite RNAs or cenRNA#4 (58% and 43%, respectively) after transfection with a scrambled sequence was similar to non-transfected cells (Supplemental Figure 4C-D). Compared to cells transfected with scrambled shRNA, we observed a decrease in the number of cells expressing centromeric RNAs (58% versus 34%) and cenRNA#4 (43% versus 22%), validating our approach of centromeric RNAs down-regulation (Supplemental Figure 4C-D). At Day 6, HeLa cells were synchronized at late mitosis - early G1 and stained for α-tubulin and CENP-B to visualize the mitotic spindle and centromeres, respectively (Figure 4A). We next scored for phenotypes related to chromosome segregation defects (Figure 4B and Supplemental Figure 5A-B). Cells transfected with scrambled shRNA displayed 30% of phenotypes related to chromosome segregation defects (Figure 4B). This percentage is explained by the high level of chromosome instability of HeLa cells (49,50). However, the down-regulation of α-satellite transcripts (α-sat) increased this percentage up to 50%, indicating the major role of these RNAs in the correct segregation of chromosomes (Figure 4B and Supplemental Figure 5B). Similarly, the proportion of cells with chromosome defects increase with cenRNA#4 down-regulation (up to 45%; Figure 4B and Supplemental Figure 5B). Overall, these data suggest that centromeric transcripts identified in these experiments are involved in faithful cell division.

## DISCUSSION

Previously, we and others have shown that transcription is required for proper centromere function in human cells and is limited to late mitosis - early G1 (19). Here, we characterize the origin, localization, and putative function of these centromere-derived long non-coding RNAs. High throughput CENP-A RIP-seq experiments revealed several novel transcripts (Figure 1), which were polyadenylated. Curiously, only 30-40% of HeLa cells had signal for cenRNAs (Figure 2). In addition, the number of foci per nucleus was also less than the number of centromeres per nucleus (Figure 2 and Figure 3). Furthermore, the α-satellite RNAs and CENP-B foci were juxta-positioned (Figure 4), results that were recapitulated for chromosome-specific cenRNAs (cenRNA#4). Finally, knocking-down a chromosome-specific cenRNA resulted in mitotic defects (Figure 4). Altogether, these data reveal cenRNAs from different chromosomes are important for genome stability.

One specific question emerges from our observations. Why are cenRNA FISH signals only found in 30-40% of the cells at early G1? We consider several possible explanations. The most plausible one is that the temporal window when cenRNAs are actively transcribed is very short, thus reducing the odds of capturing the event by bright RNA FISH, as this requires fixing the cells. Consequently, this could mean that not all centromeres produce cenRNAs at precisely the same time. Another possibility is cenRNA molecules are long lived, and not required in high abundance, allowing the deposition of CENP-A over several cell cycles. Thus, only a fraction of the centromeres may need to undergo transcription per cell cycle, potentially sporadically. A third possibility is the very low abundance and unstable nature of cenRNAs might complicate the detection of all cenRNAs, except during active transcription, both by RIP-seq or RNA FISH. Finally, a fourth possible explanation is that cenRNAs could function both *in cis* and *in trans*. In other words, cenRNAs transcribed from one centromere are also involved in function at other centromeres. This interpretation would be consistent with the observation that most human centromeres consist of very similar α-satellite sequences, thus mildly diverged sequences may not be sufficient to inhibit trans-localization or activity of cenRNAs. In this scenario, cenRNAs are either transported to other centromeres by chaperones or by diffusion of the RNA molecules. In the former case, the responsible chaperone, likely HJURP, remains to be identified. In the latter case, proximity would be a limiting factor. The closer another centromere is to the centromere producing the cenRNAs, the greater the likelihood that diffusion results in trans use of the cenRNAs. Indeed, a recent study used a novel method to analyze Hi-C data and showed that in human lymphoblastoid cells several centromeres are in close proximity, creating regulatory communities (51). Thus, our observation that there are less α-satellite RNA FISH signals than centromere signals is consistent with the possibility of trans-acting cenRNA (Figure 5).

Past reports describe the importance of centromeric transcripts in chromosome stability (10-12,19,23). We show that down-regulating centromeric transcripts using α-satellite consensus sequence shRNA decreases the level of expression of several specific cenRNAs, leading to CENP-A depletion and chromosome defects. One may argue that in a *trans* model, the down-regulation of one specific transcript should not lead to mitotic defects, as other cenRNAs will prevent the absence of the lost cenRNAs. However, the fine-tune regulation of transcription at centromeres can impede this rescue, by not re-adjusting the level of centromeric transcripts, leading to a decrease of the cenRNAs pool. Indeed, work on human artificial chromosomes suggest that the level of transcription at the centromere is balanced by post-translational modifications present at this chromatin domain, and not by cenRNAs themselves (48,52). Indeed, too much transcription is just as deleterious for CENP-A assembly as is too little. One speculative interpretation is that over-production of HAC cenRNAs might result in transmigration of those HAC-cenRNAs to other centromeres, effectively diluting CENP-A at the HAC.

Thus, in this “Goldilocks” model, overexpression of cenRNAs might result in the concentration of cenRNA exceeding its titer, followed by mislocalization of surplus cenRNA. This mislocalization might be facilitated by histone chaperones other than HJURP (53,54), resulting in ectopic deposition of CENP-A (53,55). Ectopic CENP-A nucleosomes have the intrinsic capacity to be sites of ectopic kinetochore formation, which in turn can result in dicentric chromosomes. Ectopic CENP-A has also been found in colorectal cancers and misexpression of other kinetochore components has recently been shown to correlate with poor prognosis in various cancers (55,56). It will therefore be extremely interesting to study the relationship and co-dependence between over-expressed cenRNAs and other over-expressed centromere and kinetochore proteins.

Taken together, in this follow up report, we find that multiple cenRNAs are produced from various chromosomes, ranging in size from 300-2000 bases, and these transcripts were found to be polyadenylated and localized adjacent to functional centromeres. Furthermore, despite only 30% of cells showing cenRNA signal, knockdown of chromosome-specific cenRNAs resulted in accumulated mitotic defects. Future work will be necessary to pinpoint the mechanism by which these cenRNAs assist in CENP-A loading at centromeres. We can envision 4 models by which cenRNAs can assist in CENP-A localization. First, cenRNAs might serve as true honing sequences, using an RNA-DNA hybrid mechanism to help tether HJURP/CENP-A/H4 complexes to centromeric sequences; second, cenRNAs may be structurally involved in a pre-assembly complex that stabilizes or co-folds CENPA/H4, thus acting as co-chaperone; third, cenRNAs may sequester or protect pre-assembly CENPA/H4, preventing their fortuitious association with H3 complexes, or H3-containing chromatin before they are temporally loaded onto centromeres. Finally, based on several reports of cenRNAs binding CENP-C, it is tempting to speculate that cenRNAs in a pre-assembly complex with CENP-A/H4 and HJURP only permit association of these complexes to “active” centromeres because cenRNAs have a preference for pre-existing CENP-C-chromatin bound molecules. In this final mechanism, which is speculative and will require rigorous testing, we suggest that cenRNAs may act as a genetic-epigenetic toggle, reinforcing CENP-A localization only to active centromeres, rather than just to centromeres which contain alpha satellite sequences. This model would also address a long standing mystery in the field, wherein non-homologous CENP-As from distant species, when expressed in human cells, can localize to and rescue the loss of human CENP-A (57). In the absence of a true genetic centromere in human cells, it is has been puzzling to consider how, for example, yeast CENP-A can associate with centromeres of a species removed by several million years of evolution. In this model, it can be imagined that because the chaperone HJURP may bind non-cognate CENP-As, and can presumably also bind cenRNAs arising from active centromeres, it provides a mechanism for exogenous CENP-As to localize, and rescue human CENP-A loss from centromeres. Indeed, if intra-chromosomal use of cenRNAs is the mechanism, this might have also have implications for the evolution of centromere DNAs. The centromeres of most species consist of one type of satellite DNA. One outstanding question remains how it is possible that the centromeres of all chromosomes consist of the same type of satellite DNA. It is tempting to speculate that some sort of transposition had to have occurred. For instance, there are centromere-specific transposable elements (TEs) and the centromeres of *Arabidopsis thaliana*, which are devoid of TEs, can be invaded by an LTR retrotransposon from *A. lyrata* (58). A recent study comparing centromeres from different *Zea mays* inbred lines showed that CENP-A repositioned, followed by invasion of a transposable element (59). Altogether, centromere-specific transposition events have been shown to occur in plants and furthermore the centromere-associated protein CENP-B has been domesticated from a pogo-like transposase twice (60,61). The question that remains is, beyond the transposition events on evolutionary timescales, has transposition behavior been domesticated by centromeres via trans-acting cenRNAs. These, and other fascinating evolutionary and mechanistic questions discussed above, await future investigation.

## ACKNOWLEDGEMENT

We thank members of our lab for discussion and critical feedback; Dr. Rachel O’Neill for her generous advice and expert insights on RIP-seq and identity of cenRNA sequences over the past 3 years; Drs. Tatiana S. Karpova and David A. Ball (LRBGE, NCI) for the access to microscopy facility; and Drs. Tom Misteli, Sam John, Shiv Grewal for thoughtful comments and criticism of the manuscript. We are particularly grateful to Dr. Daniel Melters for his contributions to the discussion on the evolution of centromeric RNAs and potential retrotransposons that have invaded centromeres.

This work utilized the computational resources of the NIH HPC Biowulf cluster. (http://hpc.nih.gov) The genome sequence used in this research was derived from a HeLa cell line (https://www.ncbi.nlm.nih.gov/gap). Henrietta Lacks, and the HeLa cell line that was established from her tumor cells without her knowledge or consent in 1951, have made significant contributions to scientific progress and advances in human health. We are grateful to H. Lacks, now deceased, and to her surviving family members, for permitting us access to the WGS of HeLa cells. This study was reviewed by the NIH HeLa Genome Data Access Working Group.

## FIGURE LEGENDS

**Figure 1: New RNAs associated to CENP-A chromatin identified by RIP-seq are originating from centromeres.** (A) Schematic of RIP-seq protocol. (B) Summary of downstream data analysis pipeline (described in detail in (28)). (C-E) Screenshots of read coverage from CENP-A RIP-seq experiments for each of three identified transcripts. Input and depth-normalized read depths are presented, arbitrary scale. (F) Characteristics of novel cenRNAs identified by RIP-seq. Higher-order repeat (HOR) classification for each transcript. Data include locus coordinates, size of the sequenced RNA, higher-order repeat (HOR) or closest matching, GenBank accession, current assembly array, and repeat supra-chromosomal family (SF). (G) Size distribution of 432 detected centromeric transcripts. (H) Chromosome distribution of 432 detected centromeric transcripts.

**Figure 2: cenRNAs display discrete nuclear localization.** (A) Long non-coding RNAs NEAT1 and MALAT1 and centromeric transcripts are observed by RNA FISH in early G1 synchronized HeLa cells. (B) The frequency of cells expressing NEAT1 and MALAT1 and centromeric transcripts observed in A was quantified and tabulated (N=3).

**Figure 3: cenRNAs and CENP-B domains partially co-localized.** (A) Schematic of the co-IF/RNA FISH experiment. (B & C) G1-synchronized HeLa cells are stained for CENP-B, as marker of centromeres and centromeric transcripts (α-satellite in B and cenRNA#4 in C). Cells displaying an overlap of the IF and RNA signals were counted amongst cells expressing α-satellite transcripts and cenRNA#4. No signal was observed after RNase treatment (+RNase) (N=3).

**Figure 4: The down-regulation of cenRNAs is accompanied with chromosome defects.** (A) Schematic of the down-regulation of cenRNAs followed by co-IF experiment. (B) Transfected HeLa cells with either scrambled or α-satellite or cenRNA#4 shRNA are synchronized at late mitosis – early G1 and stained for CENP-B (marker of centromeres) and α-tubulin (marker of mitotic spindle). Cells were counted and categorized for their chromosome defects (N=3).

**Figure 5: Working model.** In a *cis* model, specific cenRNAs are transcribed from one centromere and stay associated to this specific centromere, leading to chromosome defects (*e.g.,* lagging chromosomes, chromatin bridge) of this exact chromosome and the loss of CENP-A at this centromere. On the other hand, in a *trans* model, a specific cenRNAs transcribed from one centromere will act on different chromosomes. Its loss will cause genome instability on different chromosomes.

---

**Supplemental Figure 1: Description of centromeric transcripts identified by RIP-seq**. (A) Sequence logos for the most abundant motif, in each of four subsets (∼100 each) of putative centromeric transcript sequences. Motif length and significance as reported by MEME are indicated. (B-F) Pairwise percent identity matrix of 432 centromeric transcripts detected via CENP-A Rip-seq. Percent identity is displayed in shades of grey, with lighter shades indicating more similar. Hierarchical clustering was performed, which separates the transcripts into four groups (indicated at left), based on the four top most dendrogram branch points. A color code (right) indicates the chromosome of origin for each transcript. The three transcripts described in detail in this work are indicated at right.

**Supplemental Figure 2: cenRNAs are not observed after RNase treatment.** As in figure 2, RNAs were stained by RNA FISH in early G1 synchronized HeLa cells after treatment with RNase A treatment. A long exposure staining is shown to highlight the absence of RNA FISH signal (N=3).

**Supplemental Figure 3: RNAs transcribed from centromeres are poly-adenylated.** (A) Using primers designed against centromeric α-satellite, total RNAs were amplified after retro-transcription with either poly-(dT) or random hexamer primers. (B) As in A, total RNAs were retro-transcribed and amplified with primers designed against cenRNA#4. bp: base pairs; MW: molecular weight (50bp ladder); ‐RT: no RT; ‐DNA: no cDNA (N=3).

**Supplemental figure 4: cenRNAs are down-regulated by shRNA approach.** (A) Schematic of the RNA FISH and IF experiments. (B) HeLa cells are fixed and observed under microcope. H3-GFP positive cells are tabulated. (C & D) HeLa cells are stained for α-satellite transcripts (B) or cenRNA#4 (C) two days post-transfection with an shRNA constructs. Cells displaying an RNA signal were counted and tabulated (graphs on the right) (N=4 minimum).

**Supplemental figure 5: cenRNAs depletion leads to chromosome defects.** (A) Schematic of the down-regulation of cenRNAs followed by co-IF experiment. (B) Transfected HeLa cells with either scrambled or α-satellite or cenRNA#4 shRNA are synchronized at late mitosis – early G1 and stained for CENP-B (marker of centromeres) and α-tubulin (marker of mitotic spindle). Cells were counted and categorized for their chromosome defects (N=3).

**Supplemental Table 1:** List of antibodies.

| Target Protein | Reference | Manufacturer | Method |
| --- | --- | --- | --- |
| CENP-A “3-19” | ab13939 | Abcam | RNA Immuno-Precipitation – sequencing, Immuno-Fluorescence |
| CENP-B “H65” | sc-22788 | Santa Cruz | Immuno-Fluorescence |
| α-Tubulin<br>“DM1A” | 62204 | Thermo Fisher Scientific | Immuno-Fluorescence |

**Supplemental Table 2:**
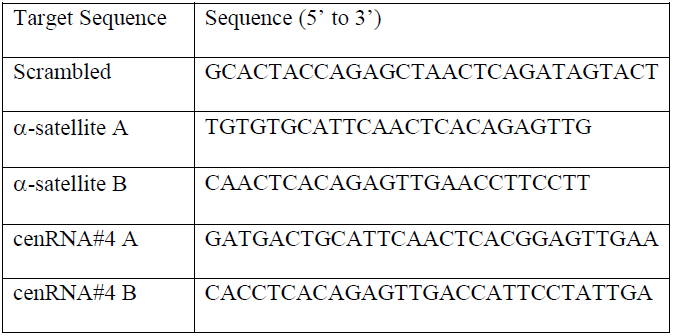
List of shRNA sequence.

**Supplemental Table 3:**
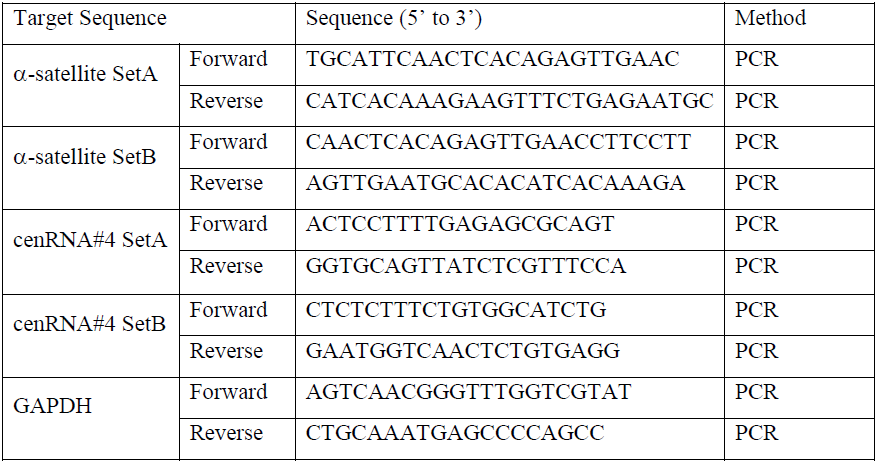
List of primer sequence.

**Supplemental Table 4.**
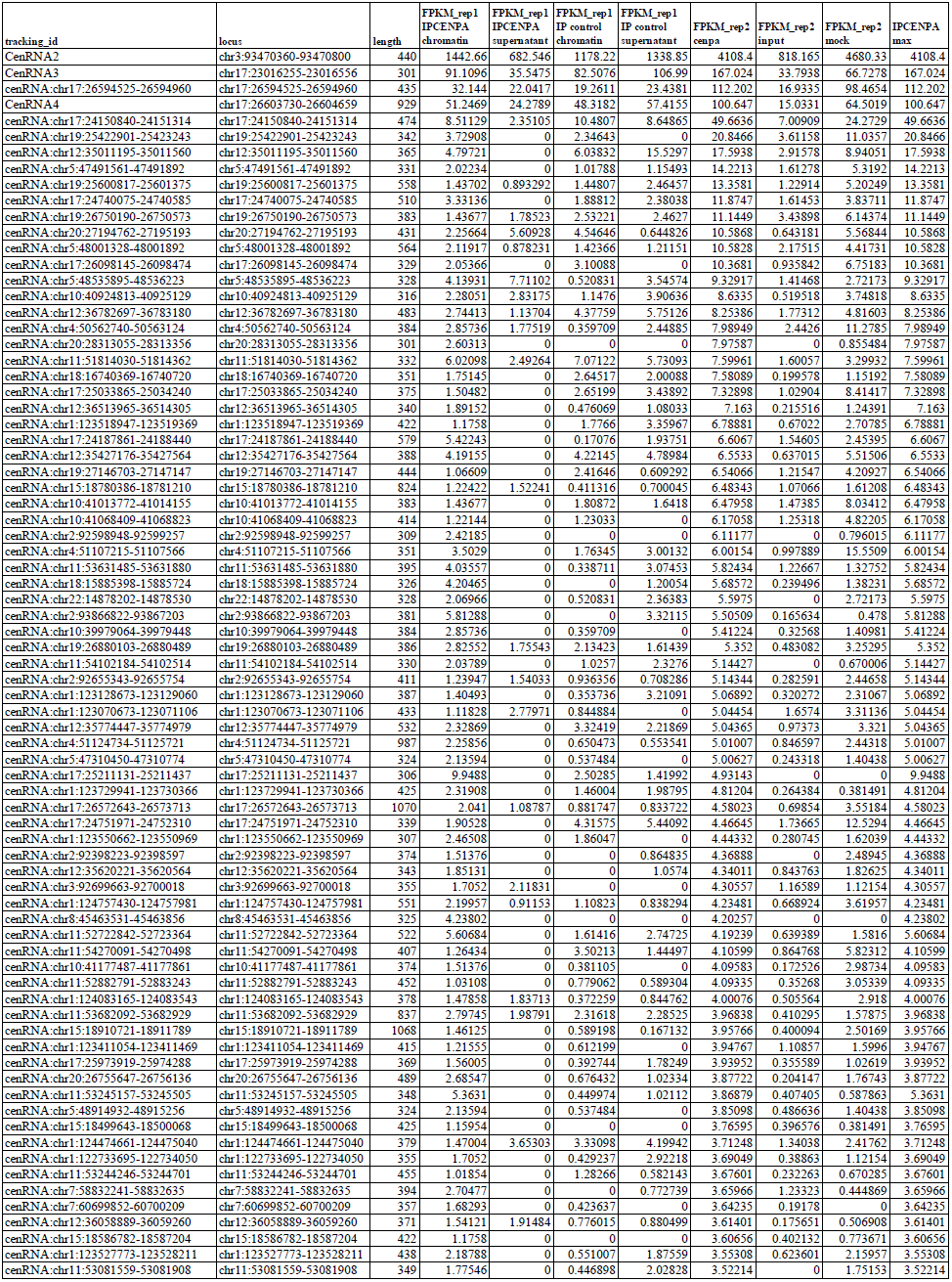
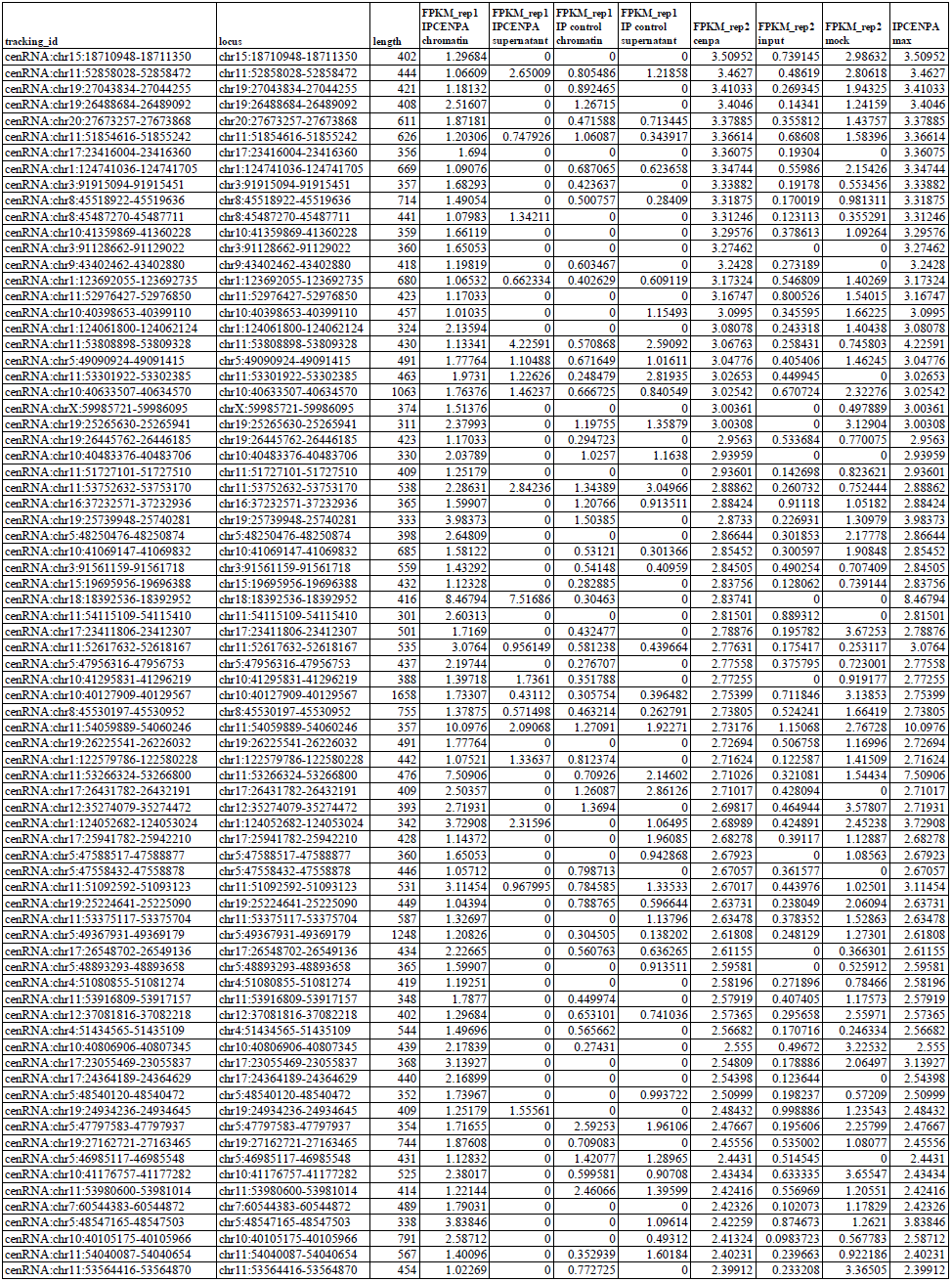
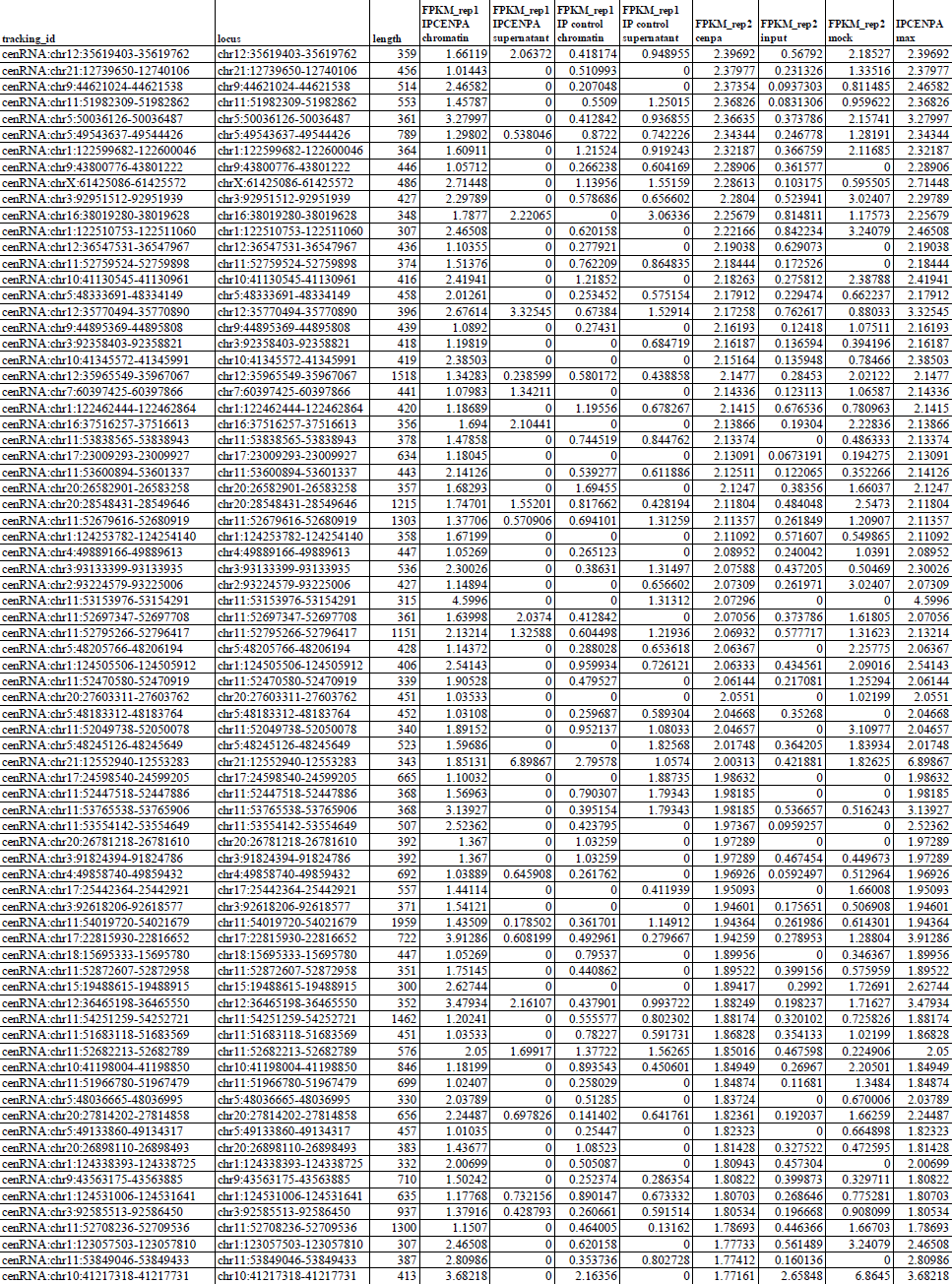
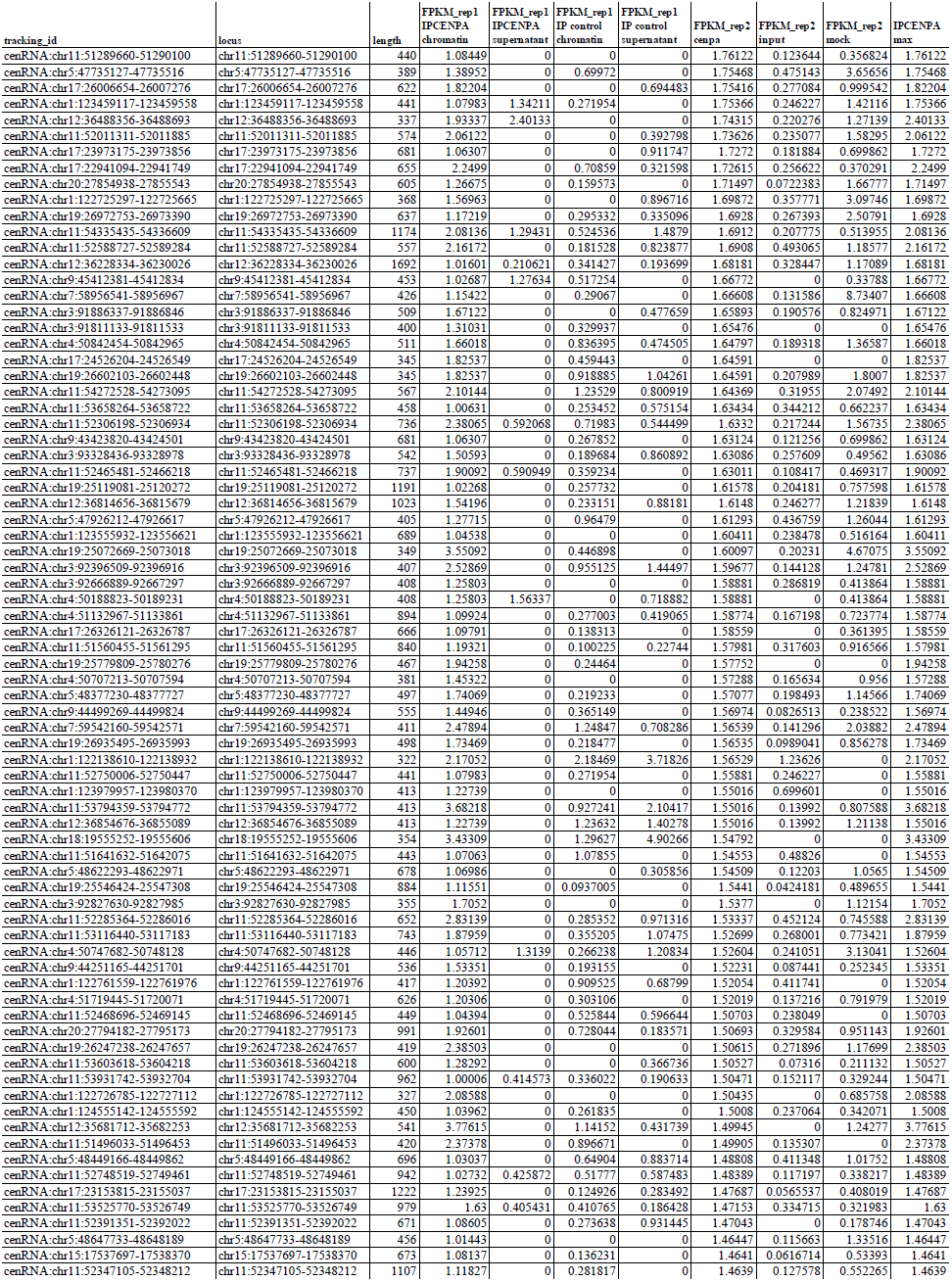
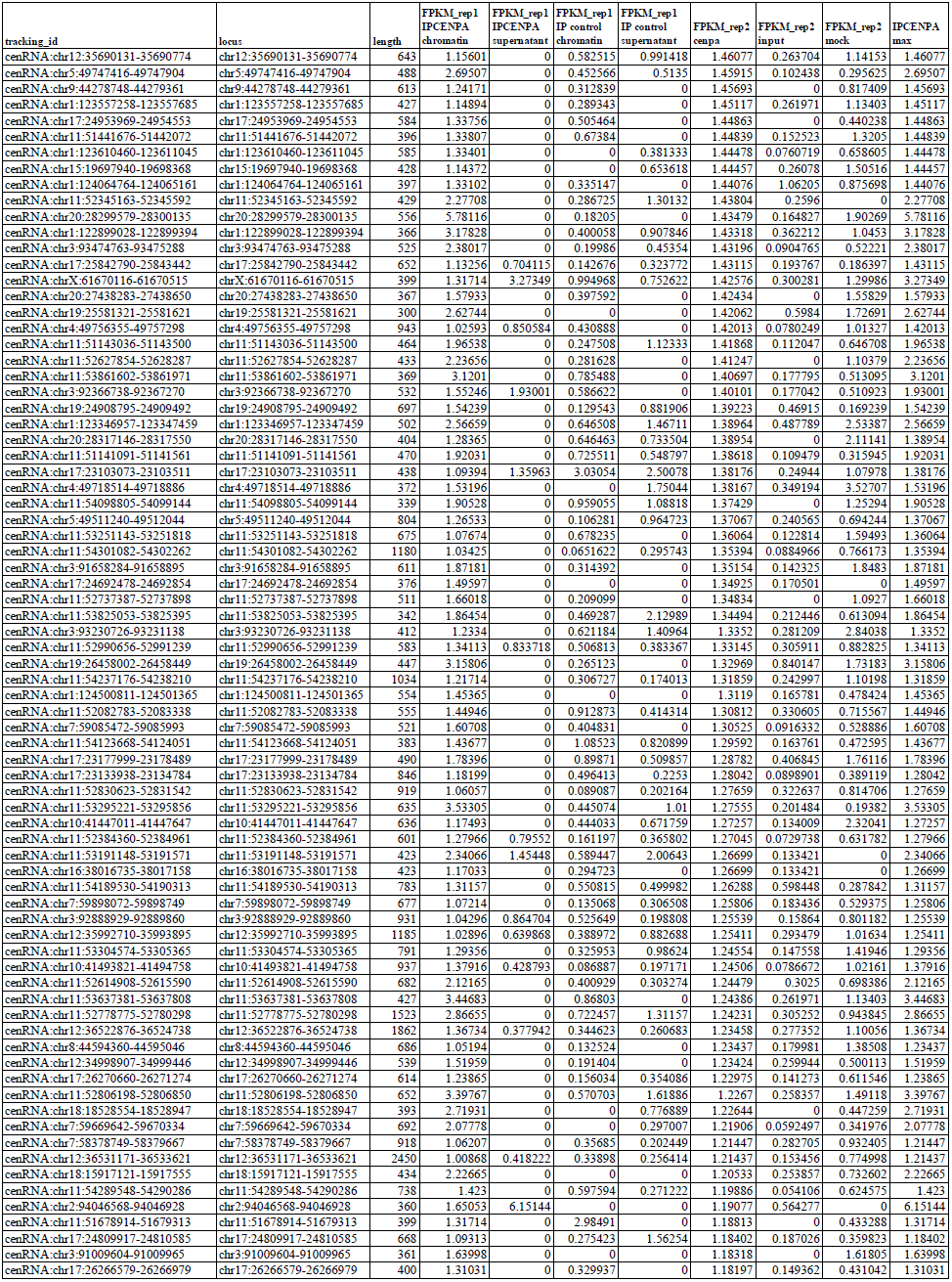
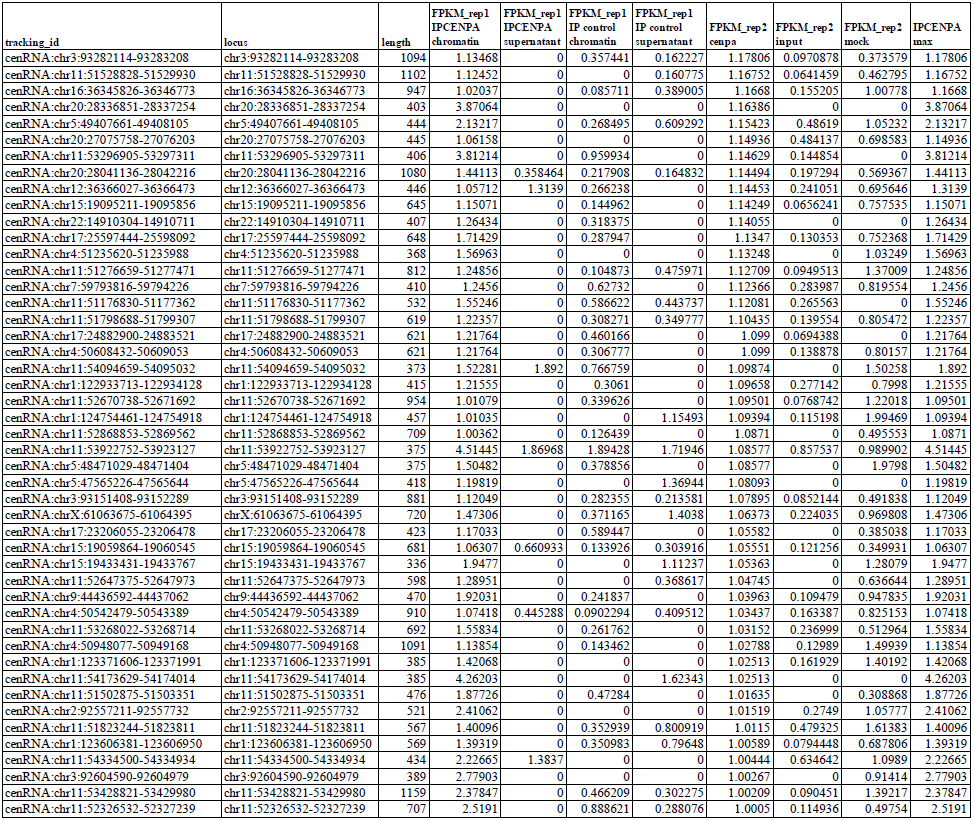

